# A biophysical limit for quorum sensing in biofilms

**DOI:** 10.1101/2020.11.01.364125

**Authors:** Avaneesh V. Narla, David Borenstein, Ned S. Wingreen

**Affiliations:** Department of Physics, University of California, San Diego, 9500 Gilman Drive, La Jolla, CA 92092; Charity Navigator, Princeton University, Princeton, NJ 08544; Department of Molecular Biology, Princeton University, Princeton, NJ 08544; Lewis-Sigler Institute for Integrative Genomics, Princeton University, Princeton, NJ 08544

**Keywords:** quorum sensing, biofilms, agent-based modeling, nutrient-limited communication

## Abstract

Bacteria grow on surfaces in complex immobile communities known as biofilms, which are composed of cells embedded in an extracellular matrix. Within biofilms, bacteria often interact with members of their own species, and cooperate or compete with members of other species via quorum sensing (QS). QS is a process by which microbes produce, secrete, and subsequently detect small molecules called autoinducers (AIs) to assess their local population density. We explore the competitive advantage of QS through agent-based simulations of a spatial model in which colony expansion via extracellular matrix production provides greater access to a limiting diffusible nutrient. We note a significant difference in results based on whether AI production is constitutive or limited by nutrient availability: If AI production is constitutive, simple QS-based matrix-production strategies can be far superior to any fixed strategy. However, if AI production is limited by nutrient availability, QS-based strategies fail to provide a significant advantage over fixed strategies. To explain this dichotomy, we derive a novel biophysical limit for the dynamic range of nutrient-limited AI concentrations in biofilms. This range is remarkably small (less than 10-fold) for the realistic case in which a growth-limiting diffusible nutrient is taken up within a narrow active growth layer. This biophysical limit implies that for QS to be most effective in biofilms, AI production should be a protected function not directly tied to metabolism.

**Significance Statement:** Biofilms are a ubiquitous form of bacterial community. Within biofilms, bacteria communicate via chemical signals in a process called quorum sensing (QS). However, if signal production is nutrient-limited, then the nutrient-deficient interior of a biofilm cannot contribute to QS, which limits the ability of bacteria to assess their own population and behave accordingly. Numerical simulations of competitions among biofilm bacteria led us to discover a novel biophysical limit on the efficacy of nutrient-limited QS. In view of this limit, we conclude that to be most effective, QS signal production should be a prized function that is not metabolically slaved.

---

Bacteria are capable of communicating with their neighbors through a process known as quorum sensing (QS). QS depends on the secretion and detection of small, diffusible molecules known as autoinducers (AIs), whose concentration increases with increasing cell density (1, 2). QS allows bacteria to control processes that are unproductive when undertaken by an individual but effective when undertaken by all members of the group, and thus leads to a competitive advantage for bacterial communities that employ QS (1–6).

QS is known to promote and regulate bacterial biofilms: immobile communities of cells densely packed in an extracellular matrix (7). QS has been demonstrated to be critical to proper biofilm formation (8–13). For example, *Pseudomonas aeruginosa* mutants that do not synthesize AIs terminate biofilm formation at an early stage (14). Given that the interior of biofilms is known to be nutrient-deficient (15), it is an open question to what extent these interior cells participate in AI production. Indeed, in many cases AI production relies on central metabolic compounds. For example, a substrate for synthesis of the ubiquitous acyl-HSL AIs is produced by one-carbon metabolism, which is highly dependent on nutrient availability (16–18). Thus, we sought to understand whether QS can be advantageous to cells in a biofilm if AI production depends on access to nutrients.

One context in which QS has been found to afford a competitive advantage in biofilms is via regulation of production of the extracellular matrix, which is composed of biopolymers, including polysaccharides, proteins, nucleic acids, and lipids. Advantages provided by the matrix include adhering cells to each other and to a substrate, creating a protective barrier against chemicals and predators, and facilitating horizontal gene transfer.

In simple models of biofilms that incorporate realistic reaction-diffusion effects, Xavier et al. (19) found that matrix production allows cells to push descendants outwards from a surface into a more O_2_-rich environment. Consequently, they found that matrix production provides a strong competitive advantage to cell lineages by suffocating neighboring non-producing cells (19). Building upon this work, Nadell et al. (20) showed that strategies that employ QS to deactivate matrix production in mature biofilms can yield a further advantage by redirecting resources into reproduction, and this scenario has been replicated and further developed (21–24). Notably, all these models assume constitutive AI production with no dependence on nutrient availability (20–24). We were therefore inspired to ask, does QS still provide an advantage in regulating matrix production if AI production is limited by nutrient availability?

To this end, we simulated competitions among biofilm-forming cells, comparing matrix-production strategies that employ QS with strategies that do not. While QS cells that constitutively produce AI could outcompete all fixed strategies, we found, surprisingly, that QS cells which produce AI in a nutrient-dependent manner have essentially no advantage over non-QS cells. We trace this result to a novel biophysical limit on the dynamic range of AI concentrations if AI production is nutrient limited. This biophysical limit applies to all bacterial systems that employ QS. These results suggest that for QS to be effective in biofilms and other conditions where nutrients are limiting, cells must privilege AI production despite the metabolic cost. From this perspective, autoinduction itself, i.e. positive feedback on AI production from AI sensing, can be viewed as one way that cells decouple AI production from metabolism.

## Results

### Agent-based Model

For simplicity and ease of visualization, we performed simulations with agent-based models (ABMs) on a two-dimensional square lattice (Fig. 1*A*). ABMs represent a system as an ensemble of autonomous agents which interact with one another according to predefined behaviors (25). In our simulations, a square can be occupied by a cell, an equivalent volume of matrix, or be unoccupied. Cells start at the bottom of the simulation domain, which is taken as the substrate to which the biofilm adheres. Cells may reproduce and form identical copies of themselves or produce matrix (see Fig. 1*B* and *C*). Matrix itself performs no actions, but fills space.

**Fig. 1.**
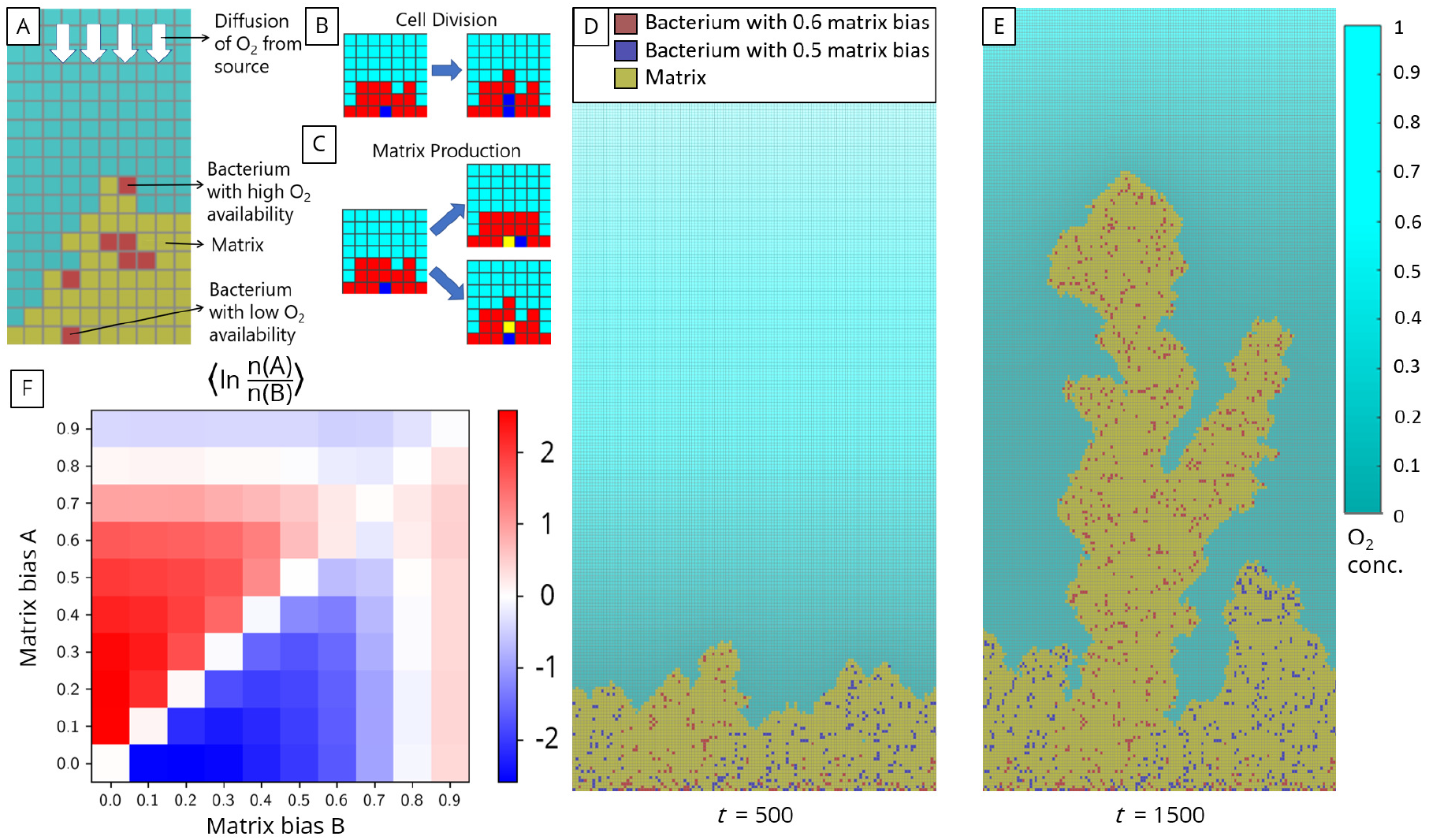
Simulated competitions of matrix-producing biofilms that grow on a submerged surface toward an O_2_ source. For details please see *Materials and Methods* and *SI Appendix*. (*A*) All simulations are performed on a 2D square lattice of width 128 sites and height 256 sites. Red squares are bacterial cells, yellow squares are extracellular matrix, and cyan squares are unoccupied. White arrows indicate diffusion of O_2_ from above. (*B*) Schematic of cell division and matrix production, shown for a blue cell surrounded by red cells. Cell division results in an identical cell being placed in an adjacent site. If no adjacent site is available, cells are shoved out of the way to make room for the new cell. Similarly, matrix production results in filling of an adjacent site with matrix. (*C*) Snapshot of a pairwise competition after 500 simulation timesteps. Red cells have matrix bias of 0.6 while blue cells have matrix bias of 0.5. Shade of cyan squares indicates normalized O_2_ concentration (normalized by the highest O_2_ concentration recorded for the entire simulation). (*D*) Snapshot of the same competition in *C* after 1,500 timesteps. (*E*) Mean of the natural logarithm of the final ratio of number of cells with matrix bias A to number of cells with matrix bias B. Between 75 and 350 simulations were performed for each competition.

Both reproduction and matrix production may require shoving to make an adjacent site available. Shoving is performed by first choosing a nearest vacant site and a shortest path to the chosen vacant site (both of which may not be unique); then, all occupants of the squares in the path are displaced along the path towards the vacant site. In our simulations, cells are assumed to be immotile and thus only move when shoved. Thus, the biofilm, composed of cells and the matrix they produce, increases in biomass and grows upward. Each simulation ends when 50% of the lattice sites become occupied or a cell reaches the top of the simulation domain.

Biomass production in biofilms requires nutrients. For example, aerobic biofilms depend on oxygen (O_2_) which usually diffuses in from a source located far away (15, 19, 26, 27). In our simulations, we consider a single limiting nutrient, taken to be O_2_, which diffuses from the top boundary of the simulation domain at a constant flux, mimicking a distant source (*SI Appendix*). We assume strong O_2_ uptake by bacterial cells to allow for a well-defined surface-growth layer within our small simulation domain. Since the timescale for the O_2_ concentration to come to a quasi-steady state (~20s for our simulation domain) is much shorter than the timescale of biomass production (~1 hour), we assume a separation of the two timescales.

Biomass production in the simulated biofilm is limited by O_2_ uptake, which we assume to be proportional to local O_2_ concentration. Thus, if the uptake of O_2_ is rapid, only cells in the upper layers of the biofilm have access to O_2_ and produce biomass. We define the fraction of O_2_ uptake used for matrix production to be the *matrix bias*. For the same amount of O_2_ taken up, a bacterium can produce a much greater volume of matrix than of new cells (we take the cost of matrix production to be 1/14 of the cost of reproduction on a per volume basis (28)).

### Bacterial Competitions

To estimate the optimal matrix bias for bacteria in our model, we performed pairwise competitions between different matrix-bias strategies (Fig. 1*D-F*). Starting with two strategies at a 1:1 ratio, we compared the cell counts of the different strategies at the end of the simulations, and found that a matrix bias of approximately 0.7 (Fig. 1*F*) performs better on average than any other constant matrix bias. Although the value of this “optimal” matrix bias depends on the simulation conditions (e.g., for a lower proportional cost of matrix, a higher matrix bias would be optimal), the non-zero value indicates that matrix production affords bacteria a fitness advantage in the presence of competition (a similar conclusion was reached by Xavier et al. (19) who used a realistic geometry for their simulations; Xavier et al. also utilized two limiting reactants, oxygen and a carbon substrate, and assumed Michaelis-Menten kinetics for their uptake by the bacteria (19, 29)).

As seen in Fig. 1*E*, after some time the cells of one strategy may overshadow their competitors and subsequently consume the entire flux of O_2_. Because after this time there is no further competition between strains, continued production of matrix by the “winning” strain would not increase access to O_2_, and could be viewed as a waste of resources. Thus switching to a low matrix bias strategy in the absence of competition could allow bacteria to increase their integrated reproductive rate. Following (19, 20), we hypothesized that bacteria could use intercellular communication (such as QS) to switch from a high matrix bias to a low matrix bias after having gained a monopoly over the nutrient and so perform better than any strategy with a fixed matrix bias.

To test this hypothesis, we incorporated QS into our simulations (Fig. 2). We performed pairwise competitions between strategies that employed QS and strategies that did not. We assumed QS bacteria constitutively produce diffusible AI, and detect local AI concentration to regulate their matrix bias. We modeled the matrix bias, *b*, of the QS bacteria as a Hill function,

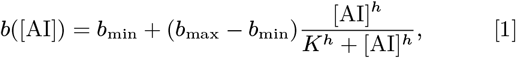

where *K* is the AI concentration at which *b* attains the value 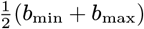, halfway between its minimum and maximum. We chose *h* = 10 to yield a near switch-like response to AI. Indeed, by varying *b*_min_, *b*_max_, and *K* we found multiple QS strategies that performed better than all fixed-matrix-bias strategies. A similar conclusion was reached by Nadell et al. (20) by employing a framework similar to Xavier et al. (19) and assuming constitutive AI production. We also tested for frequency-dependent selection effects by varying the initial seeding ratio of QS cells to fixed strategy cells (Fig. S1). While we found the QS cells to always enjoy a competitive advantage, its magnitude was reduced at very low initial QS fractions. Without the minimal number of QS cells to produce sufficient AI, these cells fail to switch from their initial high matrix bias to the low matrix bias that would allow them to capitalize on their monopoly over the diffusive nutrient. This suggests the presence of a weak Allee effect for QS cells that merits further investigation.

**Fig. 2.**
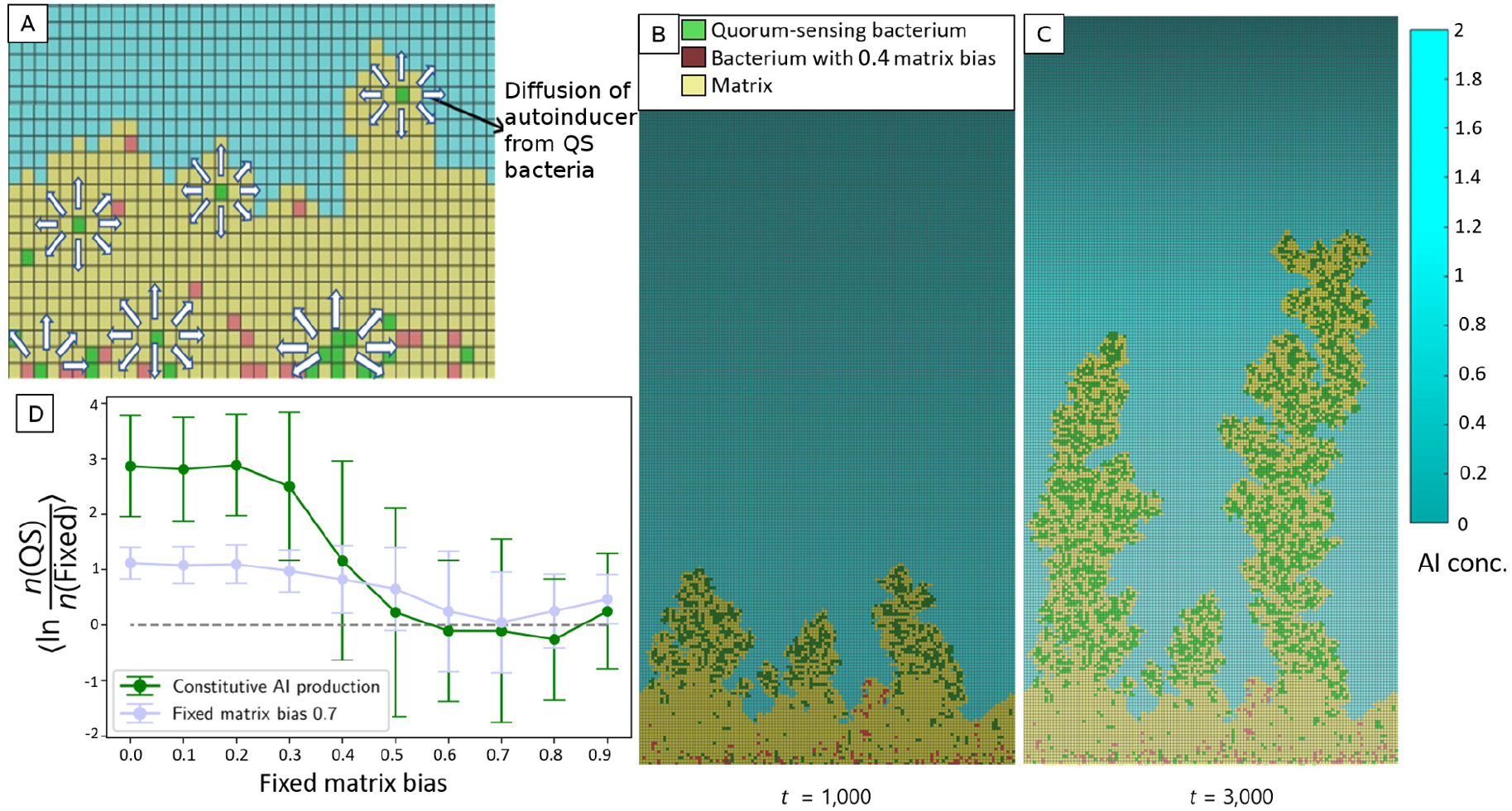
Simulated biofilm competitions between QS and non-QS cells. (*A*) QS cells shown in green produce autoinducer (AI) at a constant rate; arrows indicate AI diffusion. QS cells adjust their matrix bias based on local AI concentration. (*B*) Snapshot of a pairwise competition after 1,000 simulation timesteps. Green QS cells produce and detect AI and adjust their matrix bias from *b*_max_ = 0.9 at zero AI down to *b*_min_ = 0.1 at high AI (see Eq. 1). Non-QS cells (red) do not produce AI and have a fixed matrix bias of 0.4. O_2_ diffuses from above as in Fig. 1, but color shade now indicates local AI concentration (in arbitrary units as described in the *SI Appendix*). (*C*) Snapshot of the same competition in *B* after 3, 000 timesteps. (*D*) Mean of the natural logarithm of the final ratio of number of QS cells to number of fixed-matrix-bias cells (green curve). For comparison, the results of the pairwise competitions for the optimal fixed-strategy matrix bias of 0.7 are also shown. The error bars indicate standard deviations of log ratios. 42-65 simulations were performed for each competition.

But what if AI production is nutrient-dependent, i.e. is QS still beneficial in a nutrient-limited environment? To investigate this question, we let AI production depend linearly on local O_2_ concentration. As shown in Fig. 3, we performed pairwise competitions between fixed-matrix-bias cells and QS cells, now with nutrient-dependent AI production. Strikingly, we found that nutrient-limited QS *did not* provide a substantial competitive benefit. Specifically, nutrient-limited QS strategies had to be highly fine-tuned to ever perform better overall than fixed strategies, and at best they did not perform nearly as well as QS strategies with constitutive AI production. Notably, though the nutrient-limited QS strategies initially switched from high matrix bias to low matrix bias, the cells later switched back to a high matrix bias and thus failed to capitalize on the lack of competition.

**Fig. 3.**
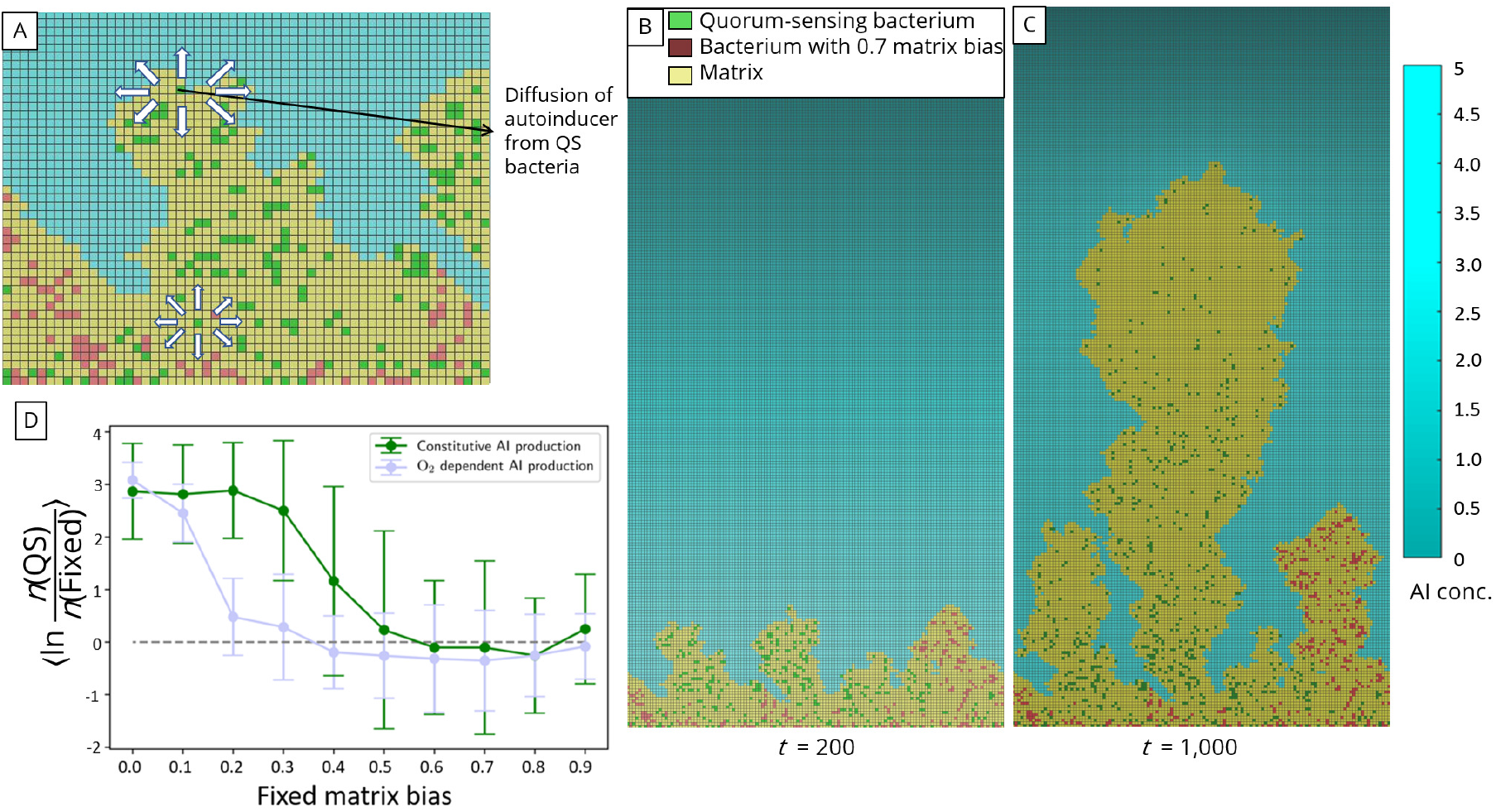
Simulated biofilm competitions between nutrient-limited QS cells that produce AI proportional to local O_2_ concentration and non-QS cells. (*A*) QS cells shown in green produce AI at a rate proportional to local O_2_ concentration (arrows highlight AI diffusion). Bacterial cells shown in red do not produce any AI. (*B*) Snapshot of a pairwise competition after 200 simulation timesteps. Green QS cells adjust their matrix bias based on local AI concentration as in Fig. 2. Red cells do not produce AI and have a fixed matrix bias of 0.7. Shade of squares indicates local AI concentration (in arbitrary units as described in the *SI Appendix*, and O_2_ diffuses from above. (*C*) Snapshot of the same competition in *B* after 1,000 timesteps. (*D*) Mean of the natural logarithm of the final ratio of number of O_2_-dependent QS cells to number of fixed-strategy cells. The error bars indicate standard deviations of log ratios. Over 60 simulations were performed for each competition.

What is it that prevents nutrient-limited QS bacteria that have achieved dominance from switching to a low matrix bias? We observed that only the cells at the edge of the biofilm produce substantial amounts of AI (as O_2_ penetration into the biofilm was designed to be low) and so the total AI production remains nearly constant. Thus, despite the increasing total population of QS bacteria, the AI concentration at the growing front of the biofilm does not increase over time. (Note that we assume a slow decay of AI, yielding a decay length of ~ 100*μ*m, to avoid artifacts associated with the finite simulation domain size.) As a result, the nutrient-limited QS bacteria are not able to distinguish between being at the edge of a large “successful” biofilm, and being part of the initial seeding density of bacteria, still in competition with other species. This contrasts with the case of constitutive AI production where the total AI production and concentration both increase with the total population of QS bacteria.

### A Novel Biophysical Limit

In our simple 2D simulations we found that nutrient-limited QS strategies provided little or no benefit to cells competing for a diffusible resource. Does this conclusion apply in more realistic settings? Perhaps surprisingly, we found that the answer is yes: There exists a corresponding biophysical limit for the efficacy of QS in 3D for bacteria whose AI production is limited by uptake of a diffusible nutrient (derivation in *SI Appendix*). Specifically, there is an upper limit on the dynamic range, DR, of possible AI concentrations experienced by cells for a given source of diffusible nutrient. For a diffusible, non-decaying AI, the minimum AI concentration, [AI]_min_, is that experienced by a single isolated cell, which senses only its own AI production. We prove that no matter how cells are arranged in 3D, the maximum AI concentration that any cell can experience has an upper bound specified by

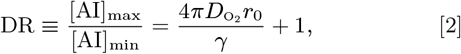

where *D*_O_2__ is the diffusion constant for O_2_ (which we take to be the limiting nutrient), *r*_0_ is the cell radius, and *γ* is the rate of intake of O_2_ per cell per concentration of O_2_. Intuitively, the biophysical limit expressed by Eq. 2 comes from recognizing that in and around a biofilm the O_2_ concentration and AI concentration are effectively mirror images (Fig. 4). This follows because O_2_ is linearly converted to AI, so local O_2_ consumption translates to local AI production, and both O_2_ and AI satisfy corresponding diffusion equations. This means that the local AI concentration can never be higher than a limit set by the minimum local O_2_ concentration, which is zero. Since a single isolated cell already experiences a finite AI concentration due to its own AI production, this upper limit on AI concentration implies an absolute upper bound on the dynamic range DR.

**Fig. 4.**
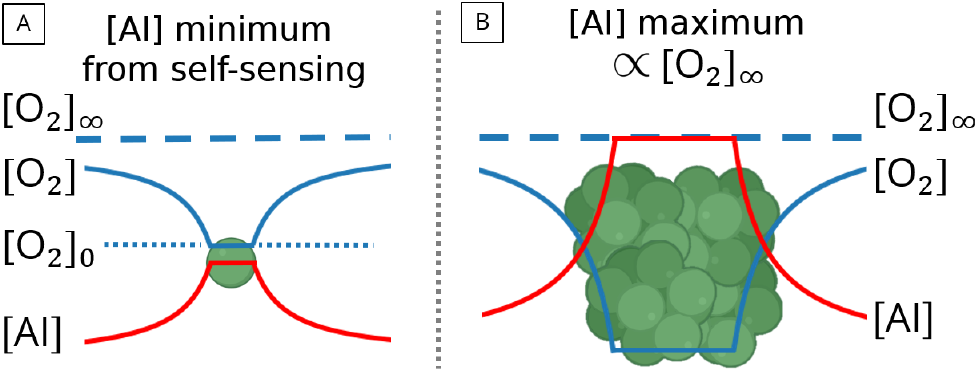
Schematic illustrating the limited dynamic range of AI concentrations. (A) A single bacterial cell, consuming oxygen and secreting AI. The AI concentration in the vicinity of the cell is proportional to the difference between the O_2_ concentration at infinity, [O_2_]_∞_, and the O_2_ concentration in the vicinity of the cell, [O_2_]_0_. (B) A bacterial colony, consuming O_2_ and secreting AI. If the O_2_ concentration inside the colony is close to zero, the AI concentration approaches a maximum value *α* [O_2_]_∞_. This relation between the AI concentration and O_2_ concentration leads to the upper limit on the dynamic range of AI described in Eqs. 2–4

Under what conditions can DR be large? Intuitively, large DR requires a small [AI]_min_ so a cell on its own must be a relatively weak producer of AI, i.e. it must be a weak consumer of O_2_. Indeed, the combination of parameters *D*_O_2__*r*_0_/*γ* in Eq. 2 is large if a single cell only weakly perturbs the local O_2_ concentration, by a combination of large values of *D*_O_2__ and *r*_0_ and a small uptake rate *γ*, which implies fast replenishment of local O_2_ by diffusion. But these conditions are not consistent with a narrow growth layer, which is precisely the case for which modeling studies have found an advantage for QS-mediated matrix production.

If AI production is metabolically slaved, what does the biophysical limit on the DR of AI concentrations imply for the efficacy of QS as a regulator of matrix production in biofilms, where growth is limited to the surface? To answer this question, note that the penetration depth of a limiting nutrient, say O_2_, into a biofilm is 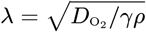, where *ρ* is the local cell density. The limit on AI dynamic range can therefore be rewritten as

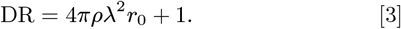

Further, the intercellular spacing is given by *ρ*^−1/3^, the number of growing layers is *n* = λ*ρ*^1/3^, and the biomass volume fraction is 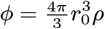. As a result, Eq. 3 can also be written as

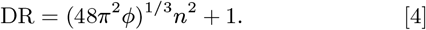

Since the volume fraction of cells near the surface of a typical biofilm is around 0.1 (30), for the limit of surface growth, i.e., *n* = 1, we obtain the DR to be less than 5. We stress that Eq. 4 is the *theoretical upper bound* for the dynamic range in such a system, and in real biological settings, the actual value may be lower.

We note that our derivation is for non-decaying AI, and the minimum AI concentration in the case of decaying AI may be arbitrary low for a cell deep in a biofilm where all AI is produced at the boundary and decays before reaching the deep interior. However, such a reduction of AI concentration is irrelevant to the collective growth strategy, since cells deep in the interior are nutrient-starved and so cannot produce substantial biomass.

## Discussion

We find that when production of a non-decaying AI is limited by a diffusible nutrient from a remote source, there exists a biophysical limit on the dynamic range of AI concentrations that cells can experience. Using agent-based simulations of biofilm growth, we demonstrate an illustrative case in which QS-based matrix-production strategies can provide a large competitive advantage – but not if AI production is limited by nutrient availability. Importantly, this biophysical limit is essentially independent of the diffusivity of the autoinducer, the size or shape of the cells, or of the concentration of the growth-limiting nutrient at its source. While our illustrative case provides a concrete mechanistic example of the effects of this biophysical limit, the limit is much more general and applies to all scenarios involving QS in nutrient-limited environments.

In principle, nutrient-limited AI production could still be exploited by bacteria in several ways: For example, in a biofilm where the density of cells is high, bacteria could employ QS to infer the concentration of the diffusible nutrient at its source. This is because, for a non-decaying AI, the local AI concentration mirrors the nutrient concentration, so that a locally-depleted nutrient with a high AI concentration would imply a large nutrient source. Further, even at lower cell densities, nutrient-limited AI could act as a single consolidated chemotactic signal that would indicate, via its negative gradient, the direction of the source of the nutrient. Alternatively, the relevant information for bacteria might be the number of *growing* cells in the vicinity rather than the total number of cells. Nevertheless, the advantages of QS seem to be much greater if AI production is not nutrient-limited.

Our results suggest a novel interpretation of autoinduction, i.e. positive feedback of AI production from AI sensing, which is a well-established feature of many QS systems (2, 31–34). It is not fully understood why autoinduction per se is desirable for cells to sense their local density. It could be presumed that a higher density of cells would necessarily result in a higher AI concentration, obviating the need for positive feedback on AI production. But this presumption would not be correct if AI production were nutrient-limited: Above a threshold cell density, AI concentration would hit its maximum as given by Eq. 2, and provide no further information. From this perspective, autoinduction may simply represent one way of breaking the dependence of AI production on nutrient availability in order to evade the biophysical limit (Eqs. 2–4, *SI Text* S1A). Another way of breaking this dependence would be for AI production to depend nonlinearly on nutrient availability, such that AI production per unit of nutrient consumed is high at low nutrient availability. In order to highlight the importance of metabolic dependence in QS, we have chosen to contrast the two extreme cases of constitutive AI production (which would be an example of such a nonlinearity) and strictly linear dependence. Real systems might fall between the two, and this is an area that merits further investigation including experimental study. In any case, either autoinduction or a nonlinear dependence of AI production on nutrients would suggest that QS is a prized metabolic function that is prioritized by the cell in nutrient-limited conditions.

Our predictions can be tested experimentally. For example, to test our prediction that AI production must be privileged, the rates of AI production in nutrient-limited and nutrient-replete conditions can be compared. If AI production is higher in nutrient-limited conditions than predicted by a strict proportionality to growth rate, it would suggest that QS is a prized function. Generally, investigation of how AI production scales with growth rate under nutrient limitation would reveal the physiological importance of QS in different growth conditions. Further, understanding the joint effect of nutrient limitation and autoinduction would reveal how the cell integrates two vital pieces of information: its local nutrient availability and the presence of other cells.

While strongly nutrient-limited AI production could be used by bacteria to infer some types of information about their environment, our main conclusion is that to reliably infer local cell density via QS, AI production should not be entirely metabolically slaved. We propose that AI production and QS are privileged bacterial functions and that despite the strong links between metabolism and AI production (16, 17, 35, 36), and the substantial cost of production of some AIs (37, 38), cells are able to decouple the two processes and regulate AI production largely independently of cell metabolism. More broadly, we hope that the results of this study will spur theoretical and experimental interest in the roles of metabolic dependency of QS signal production.

## Materials and Methods

All simulations were performed via agent-based modeling using Nanoverse (25). At each timestep, the reaction-diffusion equations for the O_2_ and AI concentrations specified by their production, consumption, and decay (if any) are solved to obtain their steady-state concentrations. This steady-state concentration determines the matrix production strategy and the probabilities in each timestep of matrix production and/or reproduction. If a cell produces matrix and/or reproduces in a timestep, then the positions of some surrounding matrix and bacterial cells are “shoved” as necessary to allow the newly produced matrix/cell to occupy a lattice site adjacent to the cell that produced it. After all matrix production and reproduction has taken place, the reaction-diffusion equations are again solved for the next timestep. This alternating procedure is repeated until the simulation halts at a pre-specified halt condition. For additional details, see *SI Appendix*.

## ACKNOWLEDGMENTS

We thank Bonnie Bassler, Farzan Beroz, Daniel Greenidge, Matthew Jemielieta, Ameya Mashruwala, Yigal Meir, and Jing Yan for stimulating discussions and input. A.V.N. acknowledges support from the Princeton University Physics Department, Office of the Dean of Research at Princeton University, UCSD Physics Department, and the qBio program at UCSD. This research was supported, in part, by National Institutes of Health Grant R01-GM082938 (to N.S.W.). This work was supported in part by the National Science Foundation, through the Center for the Physics of Biological Function (PHY-1734030). This work was done on land considered part of the ancient homelands of the Lenni-Lenape and the Kumeyaay peoples.

