## Supplemental Information for "A biophysical limit for quorum sensing in biofilms"

#### This PDF file includes:

Supplementary text
Fig. S1
Tables S1 to S3
Supplementary references

**3 Parameters** **6**

**4 Invasion Analysis** **6**

### Supporting Information Text

### Contents

### 1. Analytical derivation of biophysical limit

Here we derive the biophysical limit presented in Eq. 2 in the main text. The limit concerns the range of possible autoinducer (AI) concentrations at the surface of bacterial cells. In our model, AI production is proportional to the uptake of a diffusible nutrient (such as $O_2$). We begin by considering for simplicity a single spherical cell of radius $r_0$ centered at the origin (our results hold even if the cell is not spherical provided no dimension of the cell is much larger than another). The nutrient diffusion equation, $\frac{\partial c}{\partial t} = D_{O_2} \nabla^2 c$, becomes in steady state,

$$\frac{D_{O_2}}{r^2} \frac{\partial}{\partial r} \left( r^2 \frac{\partial c}{\partial r} \right) = 0,$$

where $c$ is the $O_2$ concentration (representing a generic diffusible nutrient) and $D_{O_2}$ is the diffusion coefficient of $O_2$ in the medium. Solving this equation with appropriate boundary conditions yields

$$c(r) = c_\infty - \frac{r_0}{r}(c_\infty - c_0), \;\; r \geq r_0, \tag{S1}$$

where we assume a constant $O_2$ concentration far from the cell, given by $c_\infty = c(\infty)$, and a fixed $O_2$ concentration, $c_0$, at the cell surface. $r_0$ is the radius of the cell. To find $c_0$, we use Fick's Law: the total rate of uptake of $O_2$ by the cell, $J_{O_2}$, is

$$J_{O_2} = D_{O_2} \oiint_S \nabla c = 4\pi D_{O_2} r_0 (c_\infty - c_0), \tag{S2}$$

where $S$ is the boundary of the cell.

What does this mean for AI production and the distribution of AI around the cell? If we suppose that the cell takes up $m$ molecules of $O_2$ in the same time that it produces one molecule of AI, the cell's rate of AI production, $J_{AI}$, is

$$J_{AI} = \frac{4\pi D_{O_2} r_0 (c_\infty - c_0)}{m} = -D_{AI} \oiint_C \nabla a(\vec{r}), \tag{S3}$$

where $D_{AI}$ is the diffusion constant for AI and $a(\vec{r})$ is the concentration of AI outside the cell.

Assuming that the cell is the only source of AI, so that the concentration of AI is spherically symmetric and zero far from the cell, and that AI decays at a rate $\beta$, we obtain

$$a(r) = \frac{D_{O_2}}{m} \frac{(c_\infty - c_0)\, r_0}{r} \exp\left( -\sqrt{\frac{\beta}{D_{AI}}}(r - r_0) \right) \left( \frac{1}{D_{AI} + \sqrt{\beta D_{AI}} r_0} \right), \;\; r \geq r_0. \tag{S4}$$

Assuming that the radius of the cell is much smaller than the length-scale of decay of AI concentration ($\sqrt{\frac{D_{\mathrm{AI}}}{\beta}}$), we can accurately approximate the AI concentration as:

$$a(r) \approx \frac{D_{\mathrm{O}_2}}{mD_{\mathrm{AI}}} \frac{(c_\infty - c_0)\; r_0}{r} \exp\left(-\sqrt{\frac{\beta}{D_{\mathrm{AI}}}}(r - r_0)\right),\;\; r \geq r_0. \qquad \text{[S5]}$$

For $r \gtrsim r_0$, we can further approximate the AI concentration as

$$a(r) \approx \frac{D_{\mathrm{O}_2}}{mD_{\mathrm{AI}}} \frac{(c_\infty - c_0)\; r_0}{r},\;\; r \geq r_0. \qquad \text{[S6]}$$

Thus, if only one cell is present, the concentration of AI at its boundary is

$$a(r_0) \approx \frac{D_{\mathrm{O}_2}}{mD_{\mathrm{AI}}}(c_\infty - c_0). \qquad \text{[S7]}$$

From Eqs. S1 and S6, we recover an approximate identity between the AI concentration and the $O_2$ at any point:

$$c(r) + \frac{mD_{\mathrm{AI}}}{D_{\mathrm{O}_2}} a(r) \approx c_\infty,\;\; r \gtrsim r_0. \qquad \text{[S8]}$$

This identity becomes exact for all $r > r_0$ in the case $\beta = 0$, when there is no decay of AI.

We now extend this result to a system of many cells. As the diffusion equations for AI and $O_2$ are linear partial differential equations in the concentration, the total distribution of the concentration values in the system is the summation of the distribution of all the cells, centered at the locations of their respective cells. Thus, by using the same boundary conditions for all cells, and for the case of $\beta \to 0$,

$$c(\vec{r}) + \frac{mD_{\mathrm{AI}}}{D_{\mathrm{O}_2}} a(\vec{r}) \approx c_\infty. \qquad \text{[S9]}$$

Above, we only require that $|\vec{r} - \vec{r_i}| \geq r_0$ where $\vec{r_i}$ is the center of any cell. Thus, the maximum AI concentration for a given set of environmental and physiological parameters is

$$\max(a(\vec{r})) = \frac{D_{\mathrm{O}_2}}{mD_{\mathrm{AI}}} c_\infty, \qquad \text{[S10]}$$

which occurs when $c(\vec{r}) = 0$. We can thus define the dynamic range $DR$, as the ratio of this maximum to the minimum AI concentration at the boundary of a cell from Eq. S8:

$$\mathrm{DR} \equiv \frac{D_{\mathrm{O}_2} c_\infty}{mD_{\mathrm{AI}}} \frac{mD_{\mathrm{AI}}}{D_{\mathrm{O}_2}(c_\infty - c_0)} = \frac{c_\infty}{c_\infty - c_0}. \qquad \text{[S11]}$$

What is the dynamic range typically obtained by bacterial cells in a biofilm? Assuming that the intake of $O_2$ is proportional to the $O_2$ concentration at the surface of the cell, the flux of $O_2$, $J_{\mathrm{O}_2}$, at the boundary of the cell is given by

$$J_{\mathrm{O}_2} = -c_0 \gamma, \qquad \text{[S12]}$$

where $\gamma$ is the per-cell rate of intake of $O_2$ (with units of $1/(\text{concentration} \cdot \text{time})$).

Equating Eq. S2 and Eq. S12, we have that

$$4\pi D_{\mathrm{O}_2} r_0 (c_\infty - c_0) = c_0 \gamma = (c_\infty - (c_\infty - c_0))\, \gamma \qquad \text{[S13]}$$

$$\implies 4\pi D_{\mathrm{O}_2} = \left(\frac{c_\infty}{c_\infty - c_0} - 1\right) \frac{\gamma}{r_0} \qquad \text{[S14]}$$

$$\implies \mathrm{DR} \equiv \frac{[\mathrm{AI}]_{\mathrm{max}}}{[\mathrm{AI}]_{\text{one cell}}} = \implies \frac{c_\infty}{c_\infty - c_0} = \frac{4\pi D_{\mathrm{O}_2} r_0}{\gamma} + 1. \qquad \text{[S15]}$$

**A. Autoinduction and metabolic regulation of AI production.** The results above presume that each bacterial cell always consumes $m$ molecules of $O_2$ for every molecule of AI that it produces, where $m$ has the same value in all conditions. However, bacterial cells may regulate their AI production so as to vary the amount of AI produced per amount of nutrient consumed. In particular, cells may increase the dynamic range of AI concentrations compared to the above limit (Eq. S15) in two ways: either by increasing the rate of AI production per molecule of $O_2$ when AI concentration is high, or by increasing the rate of AI production per molecule of $O_2$ when local $O_2$ availability is low. The former corresponds to the case of autoinduction, where the detection of AI results in activation of AI production, which is commonly observed among many bacterial species that engage in QS (1–3). The latter would require that AI production be prioritized even when nutrient availability is low and includes the case of constitutive AI production (where $m$ would be an inverse function of the limiting nutrient concentration). In either case, AI production is not metabolically slaved, i.e., AI production is not necessarily low when nutrient consumption (and subsequently metabolic activity) is low. Generally, the interior of a biofilm has both a higher AI concentration (due to the higher density of cells) and a lower nutrient concentration than the exterior. Thus, regulation of AI production in either of the forms described above would counteract the lower nutrient concentration in the interior and thus help equalize AI production per cell between the interior and the exterior of the biofilm. The effective AI production rate would then resemble the case of constitutive AI production.

### 2. Methods

**A. Agent-based modeling.** The simulations for this project were performed using Agent-Based Modeling (ABM), a simulation framework that is used widely for academic research in ecology, epidemiology, and the social sciences. It is also used in commercial and governmental uses such as business analytics, supply chain management, and military planning (4).

ABM involves representing the system of interest as a collection of autonomous actors (in our case, bacterial cells), which have well-defined behaviors and interact with one another or with the environment (4). The environment itself can be encoded as a set of variables for the simulation. The environment can also evolve with certain dynamics independent of the agents. For example, the environments in our simulations are chemicals such as nutrients used by the bacteria.

The benefit of ABM is that global dynamics emerge from many small local interactions. Only local rules are determined and communicated to the program, but one can then study the emergent global interactions.

One of the features of ABM is the representation of spatial structure. In our case, spatial structure is important as interactions among bacteria and interactions with the environment are often dictated by the spatial arrangement of cells. For example, one of the attributes we investigate is nutrient availability, where we incorporate a diffusing solute (to model nutrients such as oxygen) into the model. In such a case, spatial structure strongly influences the dynamics of the bacteria such as growth rate. For this project we used an ABM framework known as Nanoverse (4).

**B. Geometry of simulations.** The simulations were performed on a square lattice geometry in two dimensions. The shape of the complete simulation domain is a rectangle with a width of 128 squares, and a height of 256 squares. Each square in the lattice can either be empty, or be occupied by a bacterial cell, or by matrix. Each square has four neighbors. A zoomed-in region of the simulation domain (of width 7 and height 7) can be seen in Fig. 1A.

**C. Initial placement of bacteria.** At the start of each simulated competition, bacterial cells of two different strategies were placed randomly at the bottom of the simulation domain. 64 cells of each strategy were placed randomly in the squares of the bottom row, thus filling the entire bottom row.

As a test, we repeated our simulations with reduced numbers of cells at the beginning of the simulation, and found that the results did not vary substantially. Similarly, initially distributing the cells in an alternating ordered pattern did not lead to any substantial differences, and the random distribution was therefore chosen to better reflect natural conditions.

#### D. Nutrient modeling.

***D.1. Reaction-diffusion equation.*** To obtain steady-state solutions for the diffusive nutrient, we employ a discrete forward-time central-space solver to the reaction-diffusion equation (5). Though this method is valid for all diffusive nutrients with a source far from the biofilm, we refer to the nutrient in our simulations as oxygen ($O_2$). The bacterial cells are the sinks of $O_2$ in the reaction-diffusion equation. In general, the concentration of oxygen $[O_2]$ at any site is described by the following differential equation (6):

$$\frac{\partial[O_2]}{\partial t} = D_{O_2}\nabla^2[O_2] - \mu(\vec{x})[O_2]\delta(\vec{x}), \qquad [S16]$$

where $D_{O_2}$ is the diffusion constant for $O_2$, $\mu$ describes the rate of consumption of oxygen, and $\delta(x)$ is a delta function which is nonzero at each bacterial cell. This reaction-diffusion process is solved for the 2D lattice. For simplicity, the discrete grid to solve the equation is taken to be the same as the 2D lattice, i.e., grid spacing equal to the edge-length of a cell. $D_{O_2}$ is taken to be constant everywhere because even at the densest parts of the biofilm, the volume fraction of cells has been experimentally found to be less than 50% (7) and thus the value of $D_{O_2}$ in the entire domain is approximately the value of $D_{O_2}$ observed in water.

***D.2. Boundary conditions.*** To avoid edge effects, we impose periodic boundary conditions at the horizontal edges of the simulation domain. Thus, a bacterial cell that passes through the right boundary will enter from the left boundary at the same height. Similarly, the concentration of oxygen at the right edge of the simulation domain is the same as the concentration at the left edge at the same height. By imposing this boundary condition, we aim to simulate a large horizontal domain in which the biofilm can grow. Periodic boundary conditions eliminate any exceptional behavior of the biofilm at the horizontal boundaries, as the sites at the boundary are equivalent to any other sites in the domain.

***D.3. Influx of nutrient.*** A constant, spatially uniform flux of $O_2$ is introduced to the simulation domain from the top boundary. For a small simulation domain, a source of a limiting diffusible nutrient placed very far away may be approximated by a constant flux boundary condition since the amount of nutrient entering the system will be limited by diffusion and not be impacted by the details of the system (by contrast, a constant value boundary condition on the nutrient leads to a total nutrient flux that can vary with the location of the nutrient-consuming cells. The constant flux boundary condition on $O_2$ is enforced in our system by designating the top row of the simulation domain as $O_2$-producing squares that produce $O_2$ at a constant rate.

$O_2$ diffuses in the simulation environment from the top boundary towards the bacterial cells. Each bacterial cell consumes $O_2$ at a rate linearly proportional to the $O_2$ concentration at its site and acts as a sink of $O_2$. Thus, at steady state, the amount of $O_2$ consumed globally by the cells ($\delta_{\text{consumed}}$) is the same as the constant flux of $O_2$ into the whole system ($\delta_{\text{in}}$). The complete simulation domain along with the influx of $O_2$ represented by arrows (indicating the direction of flux of $O_2$) can be seen in the snapshot of a simulation in Fig. 1B.

**E. Reproduction and production of matrix.** The modeled bacterial cells can perform two actions: reproduction and production of matrix. Both reproduction and the production of matrix are stochastic processes. At every time step, the probability of a cell dividing into two cells is calculated from the formula $P(\text{reproduction}) = b_\text{r} \cdot [\text{O}_2]\Delta t$, where $b_\text{r}$ is the reproduction bias and $\Delta t$ is the duration of each time step (if the calculated probability is greater than one, the simulation is immediately halted). A random real number from a uniform distribution between 0 and 1 is then generated, and if the random number is less than the probability value for reproduction, the cell produces a copy of itself.

Similarly, the probability of matrix production is calculated from the formula $\frac{1}{2}b \cdot [\text{O}_2]\Delta t$ where $b$ is the matrix bias. Every time step, a random number is generated to decide if *two squares* of matrix will be produced. This algorithm better replicates the structure of biofilms observed in experiments than an algorithm in which squares of matrix are produced one at a time. The total biomass production rate per unit of $\text{O}_2$ is constrained such that $\chi b_\text{r} + b = 1$ where $\chi$ is the cost of reproduction relative to the cost of producing a square of matrix.

The processes of reproduction and matrix production are assumed independent: in the same time step, a cell might reproduce and also produce two units of matrix. The probability of such a double event occurring is the product of the probabilities of either event occurring, that is $\frac{1}{2}bb_\text{r}[\text{O}_2]^2(\Delta t)^2$. We chose $\Delta t$ to be small enough such that the probability of both events occurring simultaneously is very small (<0.1%).

**F. Autoinducer.** To incorporate quorum sensing, we introduce autoinducer (AI), the chemical signal produced and sensed by bacteria, as a new continuum environmental layer in Nanoverse. The sources of AI are the cells, AI diffuses, and also decays at a slow rate.. The boundary conditions for AI are periodic boundary conditions on the left and right boundaries, a reflecting (zero-flux) boundary conditions at the bottom boundary, and an absorbing boundary condition at the top boundary. An absorbing boundary condition holds the concentration at that boundary to always be 0. Thus the reaction-diffusion equation is given by,

$$\frac{\partial[\text{AI}]}{\partial t} = D_\text{AI}\nabla^2[\text{AI}] + \delta(\vec{x}) \cdot \Gamma_\text{AI}(\text{O}_2) - \beta[\text{AI}],$$

with the boundary condition that $[\text{AI}]|_\text{top boundary} = 0,\ \forall t$; $D_\text{AI}$ is the diffusion constant for AI, $\delta(x)$ is a delta function which is nonzero at the center of each AI producing cell, $\Gamma_\text{AI}$ is the rate of production of AI by that cell, and $\beta$ is the rate of decay of AI. We consider two cases: (1) constitutive AI production in which $\Gamma_\text{AI}$ is independent of local $\text{O}_2$ concentration, and (2) nutrient-limited AI production for which $\Gamma_\text{AI}$ is proportional to local $\text{O}_2$ concentration.

We note that we must assume that AI decays for our the geometry of our simulation domain. Since we have an absorbing boundary condition for the top boundary and total AI production is the same as the total $\text{O}_2$ consumption (which is constant throughout the simulation), if we don't include decay for AI, we end up with a fixed AI gradient in the region above the cells (which is equivalent to the case of a fixed flux of AI leaving the cells). Thus, when the cells line the bottom of the domain, local AI is maximal, and as the layer of actively-growing cells moves up the simulation domain, the local AI at those cells drops steadily. We include decay for AI to avoid this scenario.

**G. AI-dependent reproduction and matrix production.** In the model, The reproduction bias, $b_\text{r}$, and the matrix production bias, $b$, may depend on the local AI concentration. Specifically, $b$ is then given by the following expression:

$$b([\text{AI}]) = b_\text{min} + (b_\text{max} - b_\text{min})\frac{[\text{AI}]^h}{K^h + [\text{AI}]^h}. \qquad \text{[S17]}$$

The terms in S17 are as follows:

1. $b_\text{min}$ is the minimum possible value of $b$;
2. $b_\text{max}$ is the maximum possible value of $b$;
3. $K$ is the AI concentration at which $b$ attains the value $\frac{1}{2}(b^\text{min} + b^\text{max})$;
4. $h$ is the Hill coefficient, which characterizes cooperativity.

The matrix production bias, $b$, is still related to $b_\text{r}$ by the equation:

$$\chi b_\text{r}([\text{AI}]) + b([\text{AI}]) = 1.$$

These forms for $b$ and $b_\text{r}$ reflect the relatively switch-like behavior exhibited by many quorum-sensing bacteria. The behavior is more switch-like for higher values of $h$.

**H. Shoving algorithm.** Our simulation does not incorporate cell death, and no cells or matrix are removed from the simulation. With matrix production and reproduction, the volume of biomass strictly increases and new lattice sites become occupied. As the cells that reproduce or produce matrix may be in the interior of the biofilm, these cells must shove biomass towards the edge of the biofilm to make room for new cells/matrix. The algorithm for shoving is detailed below and the cumulative result of all steps are depicted in Figs. 1B and 1C. Steps 1-5 are common for both matrix production and reproduction. All distances are measured by $L_1$ distance, also known as Manhattan distance.

1. The closest vacant sites to the parent site (the site reproducing or producing matrix) are identified.

2. One of the closest vacant sites is chosen randomly.

3. The shortest paths to the chosen vacant site are identified. Paths are sequences of distinct lattice sites. All lattice sites in a path must be cardinal neighbors (N, S, E, or W) to the lattice sites before or after them in the path. The last lattice site in the path to a vacant site is the vacant site itself.

4. One of the shortest paths is chosen randomly.

5. All of the occupants of the lattice sites in the path are "shoved" to occupy the next lattice site in the path. Thus, the second lattice site of the path, after the active site, is now empty. This lattice site will be called the daughter site.

6. If the action is reproduction, a copy of the reproducing cell is placed in the daughter site (Fig. 1B).

7. If the action is matrix production, half of the times the matrix will occupy the daughter site (and the cell will occupy the matrix site). The other half of the time, the cell will occupy the daughter site, and the matrix will occupy the parent site. The choice between these two alternatives is made randomly.

The algorithm for shoving is extremely important as spatial structure is a crucial feature of biofilm growth. Our algorithm results in a compact biofilm as observed in nature (because the nearest vacant sites are occupied first). It also causes a homogeneous distribution of matrix and bacterial cells because the cells are pushed away from the parent site and towards the edge of the biofilm only half of the time. The other half of the time, the cells stay at the parent site.

For the bacteria and the matrix, the bottom boundary acts as a hard boundary. Thus, neither bacteria nor matrix can be shoved past the bottom of the simulation domain.

**I. Pairwise competitions.** In this investigation, we compare bacteria with different matrix production strategies and assess the competitive advantage each strategy affords in a nutrient-limited environment. To evaluate the competitive advantage of strategies, we performed pairwise competitions between bacteria of different strategies. As noted above, we started simulations with an equal number of cells of each strategy, placed randomly along the bottom row of the simulation domain. We then allowed the simulation to run until either of three specified halt conditions occurred. The first halt condition was an occupancy of 50% of the sites (that is, if 50% of the sites were filled with matrix or with bacterial cells of either strategy, the simulation would stop). The other halt conditions were any one bacterial cell reaching the top boundary of the simulation domain or the probability of either reproduction or matrix production by a bacterial cell being greater than unity.

To evaluate the competitive advantage of the strategies, we recorded the number of cells of each strategy at the end of the simulation.

### 3. Parameters

There are two relevant timescales in our simulations: the timescale of biomass production (both through reproduction and matrix production), and the timescale for the reaction-diffusion processes to equilibrate. We assume that there is a separation of timescales, as the former (typically on the order of 30-60 minutes) is much much longer than the latter (on the order of 20 seconds for a region of the size we consider). In practice, this means that we calculate the steady-state $O_2$ and AI concentrations for each fixed arrangement of cells and matrix, and then update the latter. The parameters for the reaction-diffusion equations are given in Table S1. The value of the flux of $O_2$ entering the system is essentially arbitrary, since the steady-state $O_2$ concentration simply scales with this flux, and the $O_2$ concentration is absorbed into an arbitrary linear relation between $O_2$ consumption and biomass production. Each lattice site in the domain is taken to be 1 $\mu$m by 1 $\mu$m, and this sets the length scale for the reaction-diffusion equations at 1 $\mu$m. We take the timescale for reaction-diffusion to be 1s. The diffusion constant for $O_2$ is taken to be 2000 $\mu m^2/s$, the value for $O_2$ diffusion in water at 20C at ambient pressure (8). For these parameters, the penetration depth of $O_2$ for maximally dense simulated cells is 0.3 $\mu$m. The parameters used to generate the results for quorum-sensing cells are described in Table S2.

### 4. Invasion Analysis

In principle, a particular matrix-production strategy might have a competitive advantage in a 1:1 competition with another, and yet lose that advantage when it initially constitutes a different fraction of the population. This would be an example of frequency-dependent selection. To investigate this possibility in a representative case, we performed simulations of a QS strategy versus a fixed strategy as shown in Fig. S1. We find that the constitutive QS strategy preserves its large competitive advantage when it is initially much more common than a competing fixed strategy down to when it is a few-fold lower than a competing fixed strategy. However, at very low initial fractions, the QS strategy has only a modest competitive advantage. This is because without the minimal number of cells to produce sufficient AI, the QS cells fail to switch from their initial high matrix bias to a low matrix bias that would allow them to capitalize on achieving a local monopoly on access to the diffusive nutrient.

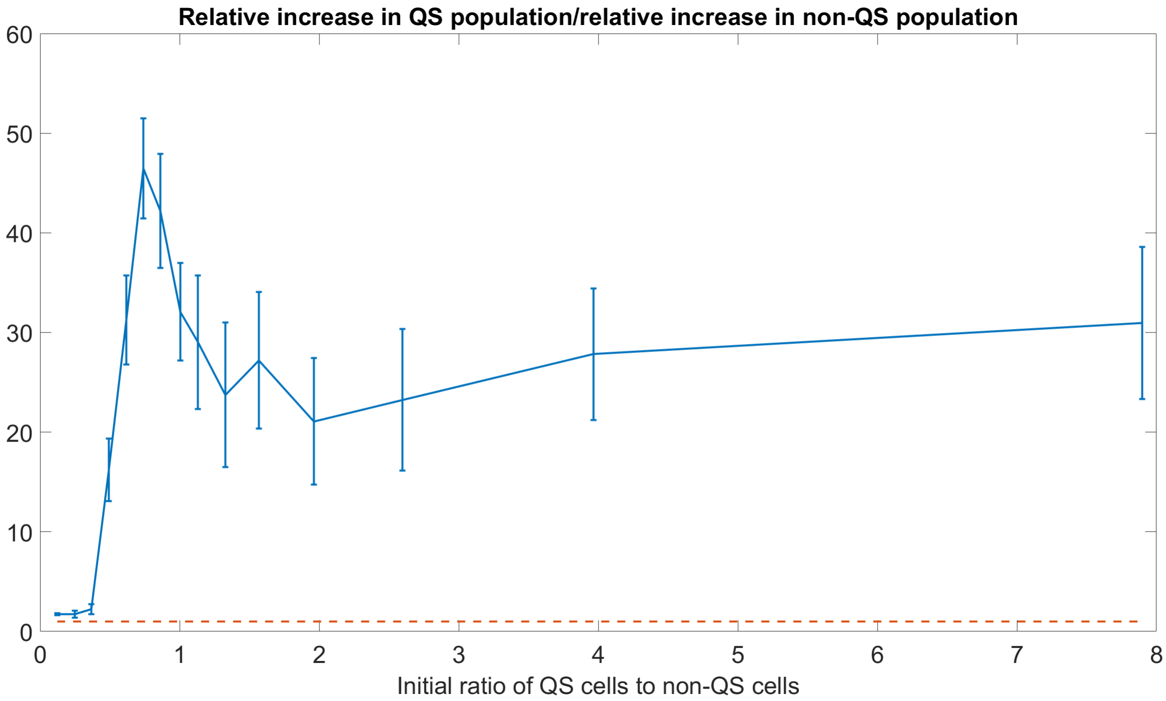

**Fig. S1.** Ratio of the increase in population of the QS strain to the increase in population of the fixed strategy strain (with a matrix bias of 0.3) for different ratios of the starting populations. The reference red line indicates ratio 1, i.e. equal performance of the two strains.

| **Property** | **Value (in dimensionless units)** |
|---|---|
| Total flux of $O_2$ entering the system | $128 \times 10^{-5}$ |
| $O_2$ diffusion constant | $2000$ |
| $O_2$ consumption rate per bacterial cell | $2 \times 10^4$ |
| Cost of reproduction relative to matrix production (for equal volumes) | 14 (9) |

**Table S1. Non-dimensionalized parameters for the reaction-diffusion equations based on units of $\mu$m and seconds.**

| Property | Description | Value |
|---|---|---|
| $h$ | Coefficient of cooperativity | $10$ |
| $b^{\text{min}}$ | Minimum value of matrix bias obtained | $0.1$ |
| $b^{\text{max}}$ | Maximum value of matrix bias obtained | $0.9$ |
| $K$ | AI concentration at which $b$ attains the value $\frac{1}{2}(b^{\text{min}} + b^{\text{max}})$ | $0.5$ |
| $\Gamma_{\text{AI}}$ | AI production rate for constitutive AI producers | $10^{-7}$ |
| $D_{\text{AI}}$ | Diffusion constant of AI | $160$ |
| $\beta$ | AI decay rate | $10^{-4}$ |

**Table S2. Parameters used to incorporate QS bacteria into the simulations in units of $\mu$m and seconds.**

| Property | Description | Value |
|---|---|---|
| $h$ | Coefficient of cooperativity | $10$ |
| $b^{\text{min}}$ | Minimum value of matrix bias obtained | $0.1$ |
| $b^{\text{max}}$ | Maximum value of matrix bias obtained | $0.9$ |
| $K$ | AI concentration at which $b$ attains the value $\frac{1}{2}(b^{\text{min}} + b^{\text{max}})$ | $3.5$ |
| $\Gamma_{\text{AI}}$ | AI production rate per unit concentration of oxygen per cell | $10^{-4}$ |
| $D_{\text{AI}}$ | Diffusion constant of AI | $160$ |
| $\beta$ | AI decay rate | $10^{-2}$ |

**Table S3. Parameters used to incorporate QS bacteria that produce AI at a rate proportional to $O_2$ uptake in units of $\mu$m and seconds.**
